# The role of conidia in the dispersal of *Ascochyta rabiei*

**DOI:** 10.1101/2020.05.12.091827

**Authors:** Ihsanul Khaliq, Joshua Fanning, Paul Melloy, Jean Galloway, Kevin Moore, Daniel Burrell, Adam H Sparks

## Abstract

*Ascochyta rabiei* asexual spores (conidia) were assumed to spread over short distances (∼10 m) in a combination of rain and strong wind. The potential distance of conidial spread was investigated in three rainfall and three sprinkler irrigation events. Chickpea trap plants were distributed at the distances of 0, 10, 25, 50 and 75 m from infected chickpea plots before scheduled irrigation and forecast rainfall events. Trap plants were transferred to a controlled temperature room (20 °C) for 48 h (100% humidity) after being exposed in the field for 2–6 days for rainfall events, and for one day for irrigation events. After a 48 h incubation period, trap plants were transferred to a glasshouse (20 °C) to allow lesion development. Lesions on all plant parts were counted after two weeks, which gave an estimate of the number of conidia released and the distance travelled. Trap plants at all distances were infected in all sprinkler irrigation and rainfall events. The highest number of lesions on trap plants were recorded closest to the infected plots – the numbers decreased as the distance from the infected plots increased. There was a positive relationship between the amount of rainfall and the number of lesions recorded. A generalised additive model was developed that efficiently described spatial patterns of conidial spread. With further development, the model can be used to predict the spread of *A. rabiei*. This is the first systematic study to show that conidia distribute *A. rabiei* over longer distances than previously reported.

## Introduction

Chickpea (*Cicer arietinum* L.) is the second most important legume crop globally and is the most widely grown legume grain crop in Australia with >1,060,000 ha harvested in 2018 (FAOSTAT 2020). Ascochyta blight caused by *Ascochyta rabiei* (syn. *Phoma rabiei*) is one of the most devastating chickpea diseases worldwide (Pande et al. 2005). With the exception of the Ord region in northern Western Australia, *A. rabiei* is the major biotic constraint to chickpea production in Australia with almost all chickpea growing areas affected (Bretag et al. 2008). *Ascochyta rabiei* survives on infected seed, volunteer chickpea plants and infested stubble, forming conidia that initiate primary infection. *Ascochyta rabiei* survives for up to two years on infested stubble at the ground surface but loses viability in up to five months when infested stubble are 5–40 cm deep in soil (W. J. Kaiser 1973; Navas-Cortés et al. 1995). Conidia are hyaline, oval to oblong in shape, none to one septate, slightly bent at one or both ends and occasionally bicelled with size range from 8.2 to 10 × 4.2 to 4.5 <u>µm</u> (Nene 1982). The optimum range for the fungus growth, pycnidial production and conidial germination is 20 °C; temperatures below 10 °C and above 30 °C have been found unfavourable for the fungus (Nene 1982; W. J. Kaiser 1973). Although there is no information on the effect of natural ultra violet light on *A. rabiei* conidia survival (Coventry 2012), exposure of cultures to wavelengths in the near ultra violet and blue range increased the rate of mycelial growth, pycnidial formation and conidial production in all isolates (W. J. Kaiser 1973).

Infected seed gives rise to infected seedlings through transmission from germinating seeds. Conidia are spread by rain splash or wind driven rain. Pycnidial formation, conidial production, host infection and disease development are favoured by temperatures between 5 and 30 °C (optimum 20 °C), relative humidity > 95 % (Nene 1982), and wetness period of 10 h or more (Khan 1999). A longer wetness period is required for spore germination at sub-optimal temperatures, and infection is rare in hot and dry conditions (Jhorar et al. 1998). Relative humidity is more important than temperature, thus *A. rabiei* development is very limited at lower humidity (e.g. 86 %) irrespective of temperature (Navas-Cortés et al. 1998). Symptoms develop within 5–6 days after infection in cool and moist (20 °C and over 90 % humidity) conditions (Khan 1999). Pycnidia develop on the infected tissues, producing conidia that are responsible for the secondary spread of the pathogen (Pande et al. 2005). Conidia are the only spore type to have been reported in Australia. Putative ascospores have been reported only once in Australia (Galloway and MacLeod 2003), however, only one mating type has been detected and detailed molecular studies have shown Australian *A. rabiei* populations to be asexually reproducing (Leo et al. 2016; Bruvo et al. 2004; Mehmood et al. 2017), thus both primary and secondary infections are attributed to conidial infection.

Rain is needed for *A. rabiei* to infect and spread within a crop; dew alone will not result in significant spread (Moore et al. 2016). During a rain event, pycnidia imbibe water and hydrostatic pressure forces conidia in the pycnidia to exude through the ostiole in a cirrhus. When a raindrop strikes the cirrhus, the kinetic energy breaks up the spore mass and disperses the conidia to nearby healthy plants, thus spreading the disease (W. Kaiser 1992; Nene and Reddy 1987; Coventry 2012). *Ascochyta rabiei* conidia have been reported to spread over 1 m by splash dispersal (Kimber 2002). The amount of kinetic energy in raindrops varies with the rainfall intensity, duration and size distribution with small drops having less kinetic energy and thus remove fewer spores (Sache 2000). Splash dispersal by large raindrops is considered more destructive because they tend to carry more spores (Sache 2000). Spores can also be washed from upper leaves to lower leaves (Sache 2000). Thunderstorms cause rapid spore dispersal from pycnidia and can result in diminution of inoculum (Coventry 2012). Low intensity, intermittent rains combined with high wind have been found to disperse spores more efficiently without diminishing the spore supply (Coventry 2012).

Irrigation is the principal mechanism for increasing grain yield, supplementing rainfall in drought seasons, or in areas receiving low rainfalls. Compared with furrow or flood irrigation, sprinkler irrigation results in the wetting of the entire crop canopy and constitutes a major source of moisture on all plant parts during dry conditions, thus increasing the risk of foliar diseases (Rotem and Palti 1969). Sprinkler irrigation plays an important role in the epidemiology of splash dispersed pathogens because spores are dispersed to their host under moist conditions and their germination is practically ensured (Lomas 1991). Unlike other forms of irrigation, water droplets generated by overhead sprinkler irrigation have the impact energy to dislodge spores (Lomas 1991). This makes it likely to play an important role in conidial spread for Ascochyta blight, *as A. rabiei* requires the impact energy of falling drops to disperse its conidia from pycnidia. Despite this, there is no study on the distance and quantity of *A. rabiei* conidia disperse during sprinkler irrigation events. Moreover, an assumption exists that conidia disperse over short distances during sprinkler irrigation events. Kimber et al. (2007) studied the spread of *A. rabiei* from primary infection foci (infected seedlings) in small plots (15 m x 9 m) in Israel. Sprinkler irrigation was used to irrigate plots, but only during dry periods, to provide conditions conducive to disease spread. Moreover, the number of sprinkler irrigation events and amount of water applied during each irrigation event were not recorded; therefore, any quantifiable relationship between irrigation, disease and the pathogen spread (via conidia) was not established.

It has been reported that *A. rabiei* conidia spread over short distances (∼1 m) via splash dispersal, and up to 10 m in the presence of rain and strong wind (in the direction of wind) (Kimber 2002; Kimber et al. 2007). However, the actual distance and number of conidia spread via wind driven rain have not been determined systematically (Coventry 2012). Kimber et al. (2007) showed Ascochyta blight spread over short distances (< 10 m) from primary infection foci (infected seedlings) in the direction of the prevailing winds. However, this was spread from a rather limited quantity of inoculum (a single infected seedling). The potential spread is likely to be greater when inoculum is plentiful. Previous studies have considered the spread of *A. rabiei* in biological rather than epidemiological terms, focusing on disease development and spread in the paddock and not on the spread of the pathogen via conidia (Coventry 2012). However, the presence of a pathogen does not automatically lead to disease development. Disease occurs due to complex interactions between a virulent pathogen, a susceptible host(s) and an environment favorable for long enough for the pathogen to cause disease (Agrios 2005).

Trapping spores on ‘trap plants’ distributed at various distances from the source of infection for a short period of time is an excellent way to investigate the distance of conidial spread. Trap plants are distributed at various distances from a source of infection for a relatively short period of time to trap spores, and then transferred to a glasshouse to allow lesion development (Salam et al. 2011). The numbers of lesions give an estimate of the number of spores released and the distance travelled. Lesion development indicates that the dispersed conidia are viable. The actual distance *A. rabiei* conidia travel from a standing crop has not been determined in such a systematic manner on a large scale. Based on preliminary work aimed to determine the distance of *A. rabiei* conidia dispersal from previous year’s infested stubble (Galloway, unpublished), we propose the pathogen is likely to spread over longer distances, than reported in the literature, in wind driven rain. Experiments conducted on plant pathogens, in a wind tunnel and rain tower, have shown that spores released by rain splash and then transported by wind are likely to be important in long-distance dispersal of pathogens (Fitt et al. 1989). Improving our understanding on how *A. rabiei* may spread in wind driven rain can help in predicting spatial patterns of conidial spread during rainfall events, and subsequent disease epidemics through modelling techniques (Coventry 2012). Models are required to predict disease risk and spread for varying environmental conditions, beyond their domain of study, as it is impractical to investigate pathogens’ epidemiology in all growing environments (Diggle et al. 2002).

To investigate *A. rabiei* conidial spread in epidemiological rather than biological terms, focusing on the pathogen spread via conidia rather than the disease development and subsequent spread in the paddock, the present study asked the following questions: (1) can conidia spread over longer distances than previously reported, and (2) do conidia spread over the same distances in rainfall and sprinkler irrigation events?

## Methods

### Study areas description

This study was conducted at three different locations, Horsham (−36.728879, 142.157311), a paddock just outside Horsham (−36.745871, 142.115969) and Curyo (−35.779312, 142.778332), in Victoria, Australia. All three sites had not had chickpea planted on them for at least 4 years. Trials established by Agriculture Victoria to assess the efficacy of fungicides against Ascochyta blight in chickpea were used to determine the distance of conidial spread in rainfall and sprinkler irrigation events. Genetically pure chickpea seed (variety PBA Striker) was sourced from the previous year’s disease free seed-increase plots. Chickpea plots were sown to dimension 10 m x 2 m on 13 May 2019. A handful (∼300 mL) of naturally infested stubble inoculum was evenly scattered across each plot at three to four nodes plant growth stage on 26 June 2019 to initiate infection. Infested residues were collected, after passing through the back of the harvester, from the previous year’s trial of similar nature. A domestic mulcher was used to break down residues into smaller pieces, and inoculum was mixed thoroughly before inoculating plots. Disease incidence for each plot was recorded as the percentage of plot with stem infections (with 0 % being no stem infections; 50 % being half the plot having stem infections; and 100 % being plot completely dead). Control plots (that received no fungicide treatments) had 90–100 % plants dead when conidia spread experiments were conducted.

### Experimental design

A method used by Salam et al. (2011) was modified for this study. At each location, 2–3 weeks old chickpea trap plants (variety PBA striker) were distributed at the distances of 0, 10, 25, 50 and 75 m from the control plots downwind of the prevailing wind direction. Downwind directions for the control plots were determined based on previous years’ weather data. Trap plants were placed at the distances of 0–75 m along radial transects (there were no other chickpea crops downwind of the control plots where chickpea trap plants were distributed). There were 10 transects separated by 10-degree increments for a total of 90 degrees between the first and the last transect. There were four trap plant stations (trap plants units) at both 0 and 10 m, and 10 trap plant stations at the distances of 25, 50 and 75 m (Table 1, Fig. S1). At the distances 0–25 m, each trap plant station consisted of 10 chickpea seedlings in two pots (five seedlings per pot). At the distances 50 and 75 m, each trap plant station consisted of three chickpea seedlings in a single pot. The distances between individual trap plant stations at each distance (0, 10, 25, 50 and 75 m) were determined using the Law of Cosine (Table 1). As there were only 4 trap plant stations at the distances of 0 and 10 m, they were placed along transects 1, 4, 7 and 10, with 30 degrees difference (space) between each trap plant stations, to increase the chances of trapping conidia. At distances of 25–75 m: all 10-trap plant stations were placed along 10 transects with 10 degrees space between the individual trap plant stations (Table 1). The locations of trap plant stations across transects were marked with flags to ensure trap plants were always exchanged in the same spot for the entire duration of the experiment (Table S1). No fungicides were applied over the duration trap plants were deployed in the paddock (Table 2).

### Spore trapping

The distance of conidial spread was determined from September through to December 2019. Spores were trapped in two rainfall events (event one ∼ 4. 6 mm, and event two ∼ 18.6 mm) in Horsham, and a single rainfall event of 0.8 mm in Curyo (Table 2). A sprinkler irrigation system (Hunter ®: Model MP300090210) located in a paddock just outside Horsham was used to determine the distance of conidial spread for three irrigation events. Sprinklers were set 110 cm above ground and were operated at 3.5 bar pressure. Each sprinkler irrigation event was equivalent to 10–12 mm of simulated rain over approximately 90 minutes (a small rain shower of ∼0.6 mm occurred during sprinkler irrigation event 3) (Table 2).

In order to determine the distance of conidial spread in rainfall and sprinkler irrigation events, trap plants were distributed in the paddock at the distances of 0–75 m from control plots before forecast rainfall and scheduled irrigation events, respectively. Trap plants were transferred to a controlled temperature room set at 20 °C for 48 hours (100 % humidity) after being exposed in the field for 2–6 days for rainfall events, and for one day for irrigation events (Table 2). The exposure period in paddock for the irrigation events was shorter for two reasons. Firstly, each irrigation event was a single continuous controlled event as opposed to natural rainfall events that fell in multiple showers over several days. Secondly, only the infected plot was being irrigated (trap plants at the distances of 10–75 m were not getting wet), therefore longer exposure period may have led to conidial desiccation on trap plants at the distances of 10–75 m. After 48 hours incubation period, the trap plants were transferred to a glasshouse (20 °C) to allow lesion development. Plants were watered daily via trays to prevent splash dispersal to nearby plants. The numbers of lesions were counted on all plant parts after two weeks giving an estimate of the number of conidia released and the distance travelled.

Weather data was recorded by an automated weather station (Measurement Engineering Australia, Adelaide, Australia). The data collected composed of date and time, air temperature (° C), dew point (° C), rainfall (mm), relative humidity (%), wind speed (ms^-1^), and average wind direction (°) recorded at 10-minutes intervals. The wind vane was not properly calibrated for the weather station at Curyo. Therefore, Birchip Cropping Group’s (BCG) agriculture farm (located 10 km away from the experimental location) weather station’s (Davis Australia) wind direction data was used for the interpretation of results (wind direction data was not included in the analysis, as wind direction is highly unstable and constantly changes). Wind direction data, recorded by Australia Bureau of Meteorology, at two locations (Hopetoun Airport 39 km, and Charlton 75 km; located in the same area as BCG and Curyo) were retrieved using the R programming language (R Core Development Team 2019) contributed packages ‘bomrang’ v0.7.0 (Sparks et al. 2017) and ‘stationaRy’ v0.5.1 (Iannone 2020) and compared with the BCG’s weather station data to ensure the BCG’s weather station wind direction data was valid (Fig. S2).

### Development of a model to describe spatial patterns of conidial spread

A preliminary linear mixed model analysis was conducted in the R programming language, using the lmer() function from the contributed package ‘lme4’ v1.1-1.23 (Bates et al. 2015), to investigate the influence of ‘spread event’ (as a random variable: any spread event regardless of sprinkler irrigation or rainfall events) and ‘experimental location’ on the number of lesions recorded. A reductive approach was used to investigate the significance of the random variables ‘spread event’ and ‘experimental location’ before modelling spatial patterns of conidial spread. Preliminary analysis showed the influence of ‘experimental location’ to be non-significant implying conidial spread patterns were consistent between experimental locations. This result allowed generalised additive model (GAM) to compare the influence of weather variables without the need to include experimental location as a dependant factor. Generalised additive models were used because they have been reported to efficiently describe the effect of various environmental variables on plant pathogen interactions (Sparks et al. 2011; Yee and Mitchell 1991). The GAM for conidial dispersion, based on the pooled data across all experimental locations for all six spread events, was fitted using the R programming language via the gam() function from the contributed package ‘mgcv’ v1.8-31 (Wood 2011). Standard information criterion, such as Akaike’s Information Criterion (AIC) (Akaike 1974), the Bayesian Information Criterion (BIC) (Gideon 1978), as well as adjusted R-squared values were compared across models to select the best fitting model.

Letting the mean number of lesions for pot, (pooled across all experimental locations for all six spread events) be the response variable *Y*_*i*_, and *x*_*i*_ be the associated covariate vector with the distance of pot, from the infected plot being *x*_*i*l_, the mean wind speed being *x*_*i*2_ and the amount of irrigation or rainfall being *x*_*i*3_, during spread events, the GAM model that achieved best fit was of the form:

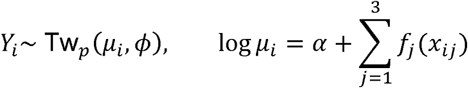

where the *f*_*j*_ are independent “thin-plate splines” which are the default smoother used by the gam() function. For details about the mathematical specification of these and other smoothing splines, see Wood (2017). The use of the Tweedie distribution family to model the response data is now standard in the case of interval scale measurements with substantial zero-inflation, i.e., a positive probability mass at zero will indicate no lesions were observed (Dunn 2004; Hasan and Dunn 2010). As it is implemented in the tw() function passed to the family parameter in the gam() function, the power parameter *p* – which relates the Tweedie variance to its mean – is estimated from the data (Wood 2011). The chosen link function for this family was the logarithm link; the canonical or natural link function for Tweedie distributions, and the default supplied to the gam() function. The gam() function also allowed the dimension of the basis for smoothed terms to be manually selected via the parameter *k*, which controls the “wiggliness” of the resultant curve. The value of *k* was chosen to be 5 (*i*,.*e*.*k* = 5) for all three predictors due to rather limited data.

### Data cleaning and visualisation

Data were processed and visualised in R using the contributed packages ‘tidyverse’ v1.3.0 (Wickham et al. 2019), ‘lubridate’ v1.7.8 (Grolemund and Wickham 2011) and ‘clifro’ (Seers and Shears 2015) for wind roses. The best fitting GAM was visualised using a custom function created for this work that uses the contributed package ‘mgcViz’ v0.1.6 (Fasiolo et al. 2019). Mean wind speed was calculated using a circular averaging function from ‘SDMTools’ v1.1-221.2 (VanDerWal et al. 2014).

## Results

### Model fit

The linear mixed model analysis indicated that experimental location had no significant effect on the number of conidia trapped and distance travelled during each spread event, thus experimental location was not used when fitting the GAMs. Thirteen GAM models were evaluated to describe spatial patterns of conidial spread. The model selected as the best fitting had an adjusted R-squared value of 0.674, AIC value of 663 and a BIC value of 709. The smooth terms for distance (D) and precipitation (P) were significant terms (p < 0.05). The smooth term for wind speed (W) was not significant. However, including this term only as a linear predictor reduced the fitness of the model, R-squared value of 0.657, AIC value of 689 and BIC value of 729, so the smoothed term was included in the final model (Fig. 1).

**Fig. 1.**
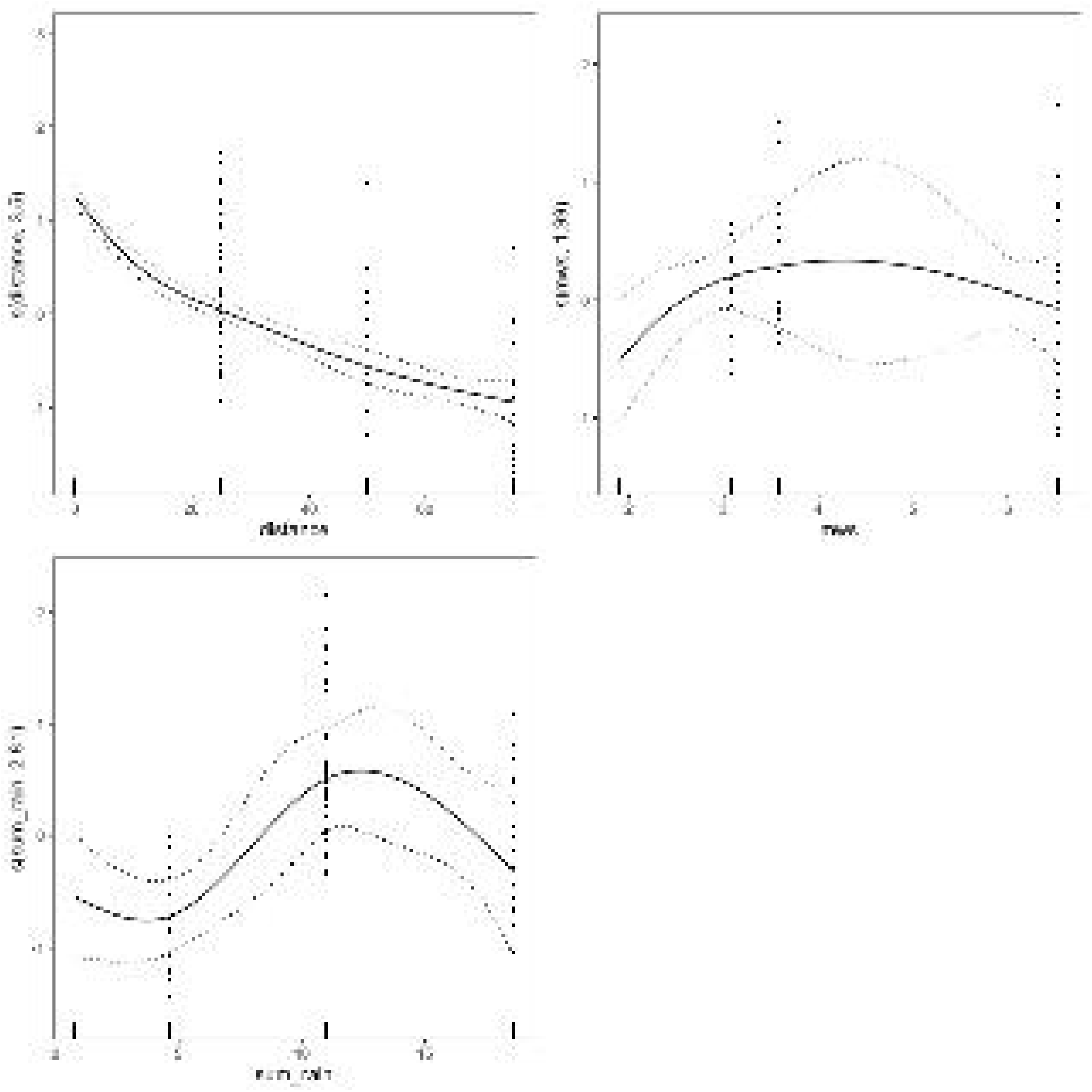
Generalised additive model fit for smoothed terms of three predictors: distance, mean windspeed (mws) and total precipitation (sum_rain)

### Dispersal in rainfall events

Trap plants at all distances (0–75 m) were infected in all 3-rainfall events, indicating conidia spread to 75 m in each rainfall event. There was a strong relationship (R^2^ = 0.657) between mean wind speed, rainfall intensity, distance from the infected plot, and the number of conidia trapped (counted as lesions). The highest number of lesions on trap plants were recorded closest to the control plot at 0 m in each rainfall event - the numbers decreased as the distance from the control plots increased (Fig. 2). The number of lesions recorded increased with increasing rainfall. At Horsham, a total of 159 lesions were recorded at all distances in rainfall event 1 (∼ 4.8 mm), and 275 lesions were recorded in rainfall event 2 (∼18.6 mm) (Table 2). In rainfall event 1 (Horsham Rain 1), a uniform conidial spread pattern across transects was observed at the distances of 0–25 m (i.e., all trap plant stations at the distances of 0–25 m were infected), whereas a non-uniform conidial spread pattern across transects was observed at the distances of 50 and 75 m (Fig. 3). Uniform conidial spread pattern across transects at all distances was observed in rainfall event 2 (Horsham Rain 2). Mean wind speed and direction varied during rainfall events 1 and 2 (Fig. 4). Mean wind direction was predominantly in the direction of trap plants (NE, SE) during both spread events (Fig. 4)

**Fig. 2.**
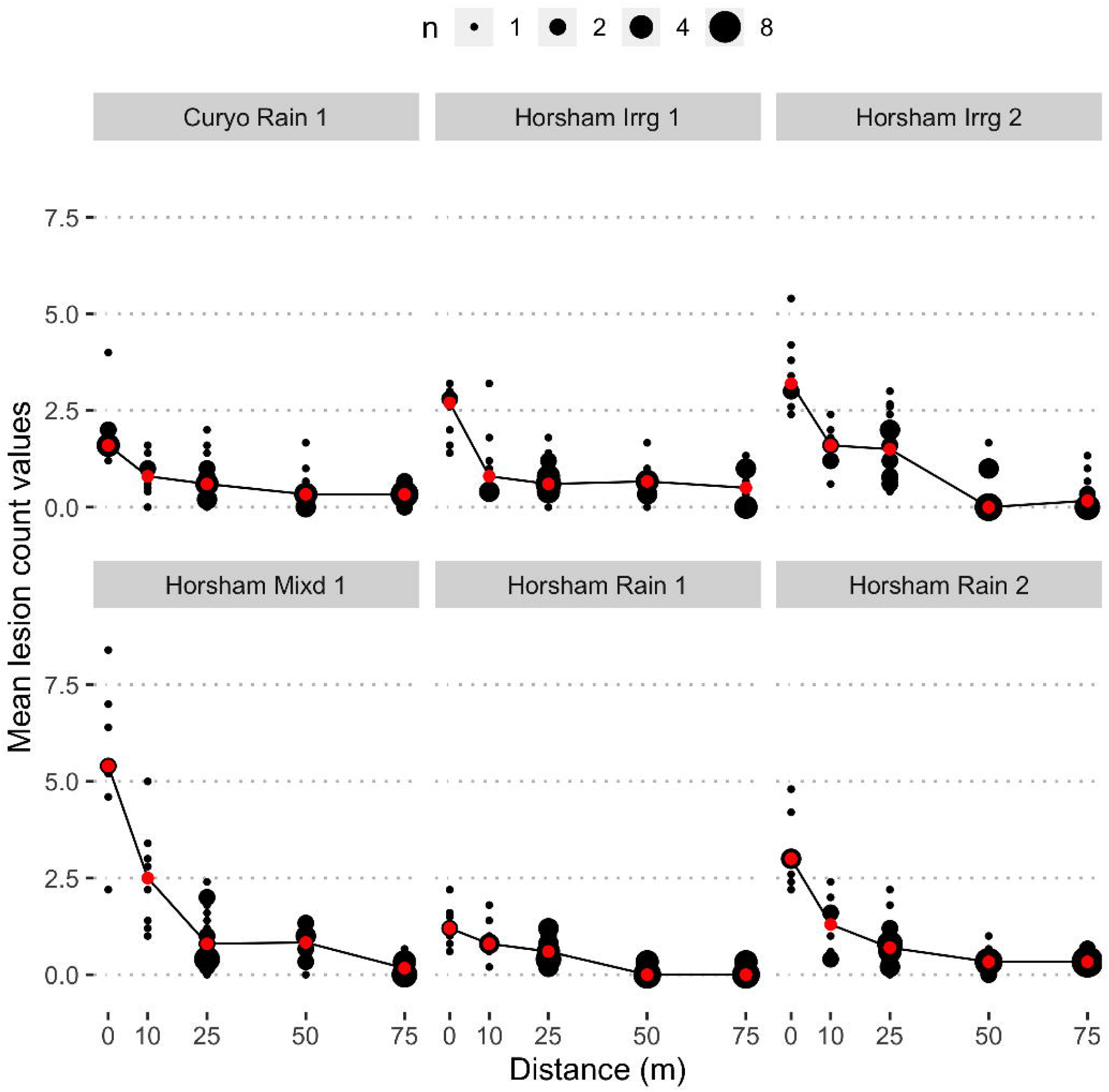
Mean number of conidia dispersed (counted as lesions) during each spread event at each distance, where ‘n’ (indicated by the point size) represents the number of pots that shared the same mean number of lesions counted per trap plant per pot. The mean count values are shown on the y-axis and distance conidia dispersed is shown on the x-axis. Red dots indicate median lesions counted per trap plant. Conidia dispersed 75 m in all irrigation and rainfall event

**Fig. 3.**
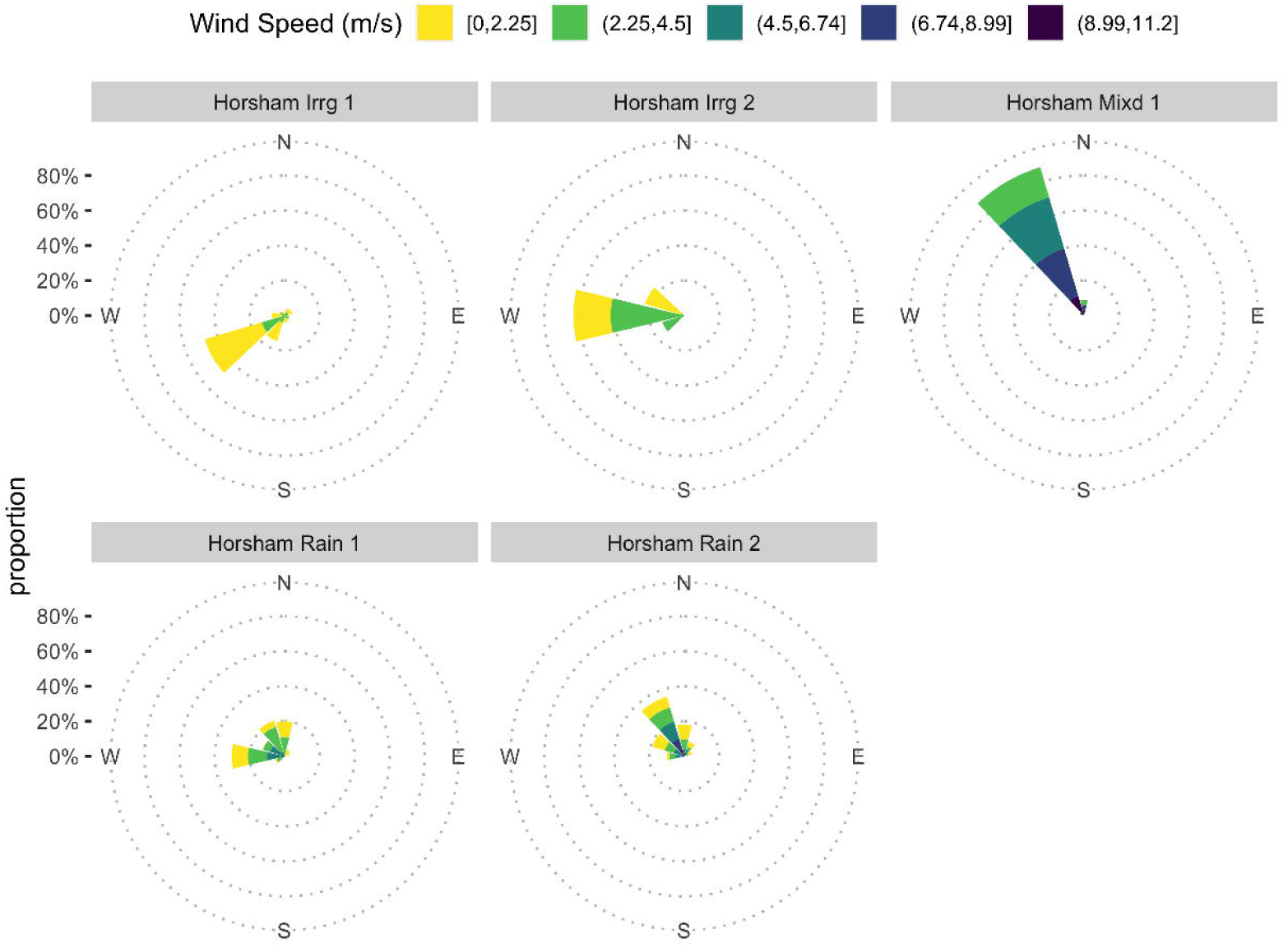
Heatmaps showing spatial patterns of conidial spread during each spread event. Spatial spread patterns were determined by counting lesions on trap plants (placed along radial transects at the distances of 0–75 m from infected plots), where ‘n’ represents mean number of lesions per trap plant per pot. Concentric circles show distance of trap plants from the infected plots: the inner single solid dot represents 0 m distance and the outer most dotted line represents 75 m distance from the infected plot

**Fig. 4.**
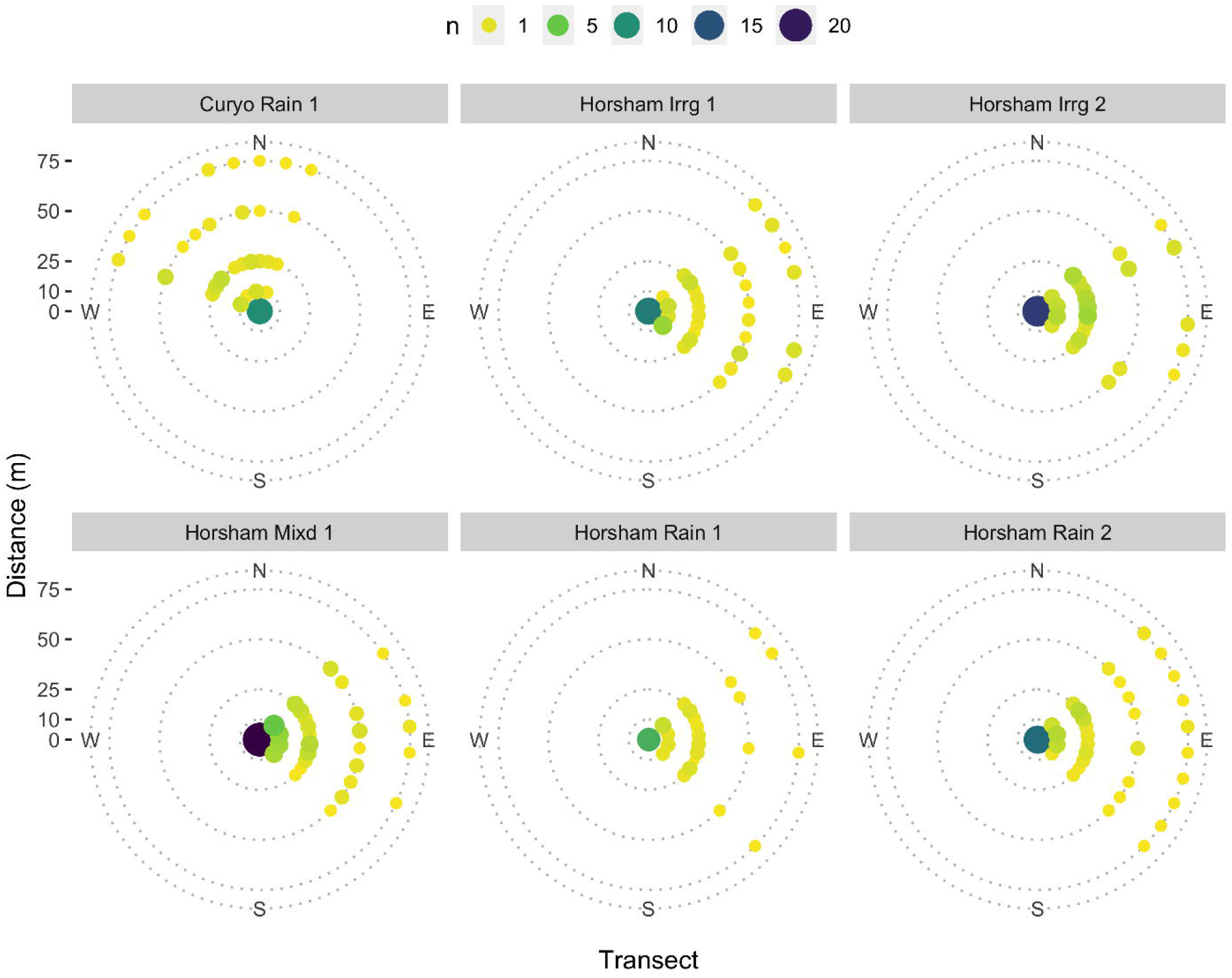
Wind roses, for all spread events except Curyo Rain 1, showing wind speed and direction recorded at 10 minutes interval

At Curyo, a total of 193 lesions were recorded in a single rainfall event (Curyo Rain 1) of 0.8 mm (Table 2). Conidial spread pattern was almost uniform across transects at the distances of 0–50 m, while a non-uniform conidial spread pattern across transects was observed at 75 m (Fig. 3). Mean wind speed and direction varied during the spread event (Fig. S2). Mean wind direction was not in the direction of trap plants (Fig. S2) but, compared to rainfall events-1 and 2, wind direction appeared to have limited influence on the number of lesions recorded (Table 2), and the spatial patterns of conidial spread at Curyo (Fig. 3).

### Dispersal with sprinkler irrigation

Trap plants at all distances (0–75 m) were infected in all three irrigation events. There was a strong correlation between mean wind speed, irrigation, distance from the infested plot and the number of conidia trapped (counted as lesions). The highest numbers of lesions on trap plants were recorded closest to the control plot in each irrigation event - the numbers decreased as the distance from the control plot increased (Fig. 2). The total number of lesions recorded on trap plants varied greatly between all three irrigation events, with a total of 250 lesions recorded in event 1 (∼10–12 mm), 368 lesions in event 2 (∼10–12 mm) and 477 lesions in event 3 (∼10–12 mm irrigation event and a small rain shower of 0.6 mm) (Table 2). In event 1 and 3 (Horsham Irrg1 and Horsham Mxd1), a uniform conidial spread pattern across almost all transects was observed at the distances of 0–50 m, whereas conidial spread pattern across transects was not uniform at 75 m (Fig. 3). In event 2: conidial spread pattern across transects was uniform at the distances of 0–25 m, whereas a non-uniform conidial spread pattern across transects was observed at the distances of 50 and 75 m (Fig. 3). Mean wind speed and direction varied between each spread event (Fig. 4).

Maximum mean wind speed (6.5 ms^-1^) (Table 2) and maximum gust (11.2 ms^-1^) were recorded was recorded during irrigation event 3. Mean wind direction was predominantly in the direction of trap plants (NE, SE) during all spread events (Fig. 4).

## Discussion

Conidia were shown to distribute *A. rabiei* over longer distances than previously reported. Conidia dispersed 75 m from infected plots in all sprinkler irrigation and rainfall events. The mean number of lesions recorded on trap plants decreased as the distance from infected plots increased. There was a strong relationship (R^2^ = 0.657) between mean wind speed, the amount of precipitation (rainfall and/or sprinkler irrigation) and the number of conidia trapped. The experimental location had no significant effect on the number of conidia trapped and distance travelled during each spread event (P>0.01). A model was developed that efficiently described spatial patterns of conidial spread under field conditions. Given more data and further development, this model can be used to predict the spread of *A. rabiei*.

The highest mean number of lesions on trap plants were recorded closest to the control plots in all spread events – the numbers decreased as the distance from the control plots increased. These results support the findings from previous studies (Coventry 2012; Kimber et al. 2007) who showed *A. rabiei* inoculum density decreased as the distance from infected foci increased. A similar dispersal pattern has been observed for other splash dispersed pathogens, such as *Colletotrichum acutatum* (Yang et al. 1990), *Parastagonospora nodorum* (previously known as *Septoria nodorum*) (Griffiths and Ao 1976), and *Ascochyta fabae* (Pedersen et al. 1994). For splash dispersed pathogens, plants close to the source of infection are more exposed to spore load due to localised splash dispersal and become infected quickly, resulting in more lesions (Coventry 2012). More lesions were recorded on trap plants with increased rainfall and in an event with a combination of a small rain shower and a sprinkler irrigation event compared to a sprinkler irrigation event alone (in case of sprinkler irrigation event 3, i.e., Horsham Mixd), suggesting that increased precipitation dislodged more conidia, which were dispersed by wind driven rain. The amount of rainfall has been reported to have a major influence on Ascochyta blight severity and the distance disease spreads (Coventry 2012).

Wind speed and direction coinciding with increased rainfall influenced the spatial patterns of conidial spread. Uniform conidial spread across transects at all distances was observed in rainfall event 2 (Horsham Rain 2), which indicates strong winds from NW directions coincided with increased rainfall and dispersed conidia across all transects downwind (SE, NE) from the infected plot. Although trap plants were located downwind in rainfall event 1 (Horsham Rain 1), uniform conidial spread across all transects was not observed at 50–75 m, which reflects the influence of reduced rainfall and rather low wind speed (compared to rainfall event 2) during the spread event. Compared to rainfall event 1 at Horsham, the amount of rainfall and wind direction appeared to have limited influence on the number of conidia trapped and the spatial patterns of conidial spread for rainfall event 3 (Curyo Rain 1) at Curyo. More lesions (193 lesions) were recorded in a single rainfall event of 0.8 mm at Curyo, although mean wind direction was not in the direction of trap plants. In addition, conidial spread pattern was almost uniform at the distances of 0–50 m. Conversely, fewer lesions (159) were recorded in a rainfall event of 4.8 mm at Horsham when trap plants were located downwind; and conidial spread pattern was uniform at the distances of 0–25 m only. However, a direct comparison between these two rainfall events is not possible due to a location difference and rather limited data. Furthermore, our model does not consider continuously changing wind direction and dynamics associated with turbulence, which may assist in explaining variation in spatial patterns of conidial spread. Conidia dispersed 75 m and spatial pattern of conidial spread was uniform across transects at the distances of 0–50 m, when the amount of rainfall was as low as 0.8 mm (Curyo Rain 1 event with mean wind speed of 3.6 ms^-1^); and mean wind direction was not predominantly in the direction of trap plants. This suggests that only a small amount of rain is required to dislodge a large number of conidia. A similar dispersal pattern has been reported for the sexual spores (ascospores) of *A. rabiei*, where ascospores discharge depended more on the occurrence of rain than the amount (Trapero-Casas et al. 1996).

More conidia (368) were trapped in sprinkler irrigation event 2, compared to sprinkler irrigation event 1, but conidial spread pattern across transects was not uniform at the distances of 50–75 m. Conversely, fewer conidia (250) were trapped in sprinkler irrigation event 1, but conidial spread pattern was almost uniform across transects at all distances. The lower number of spores and non-uniform spatial spread pattern for sprinkler irrigation event 2 is rather surprising because an equal amount (∼10–12 mm) of irrigation was applied in both sprinkler irrigation events. Moreover, stronger mean wind speed was recorded for sprinkler irrigation event 2 and the non-infected trap plants were located downwind (the mean wind direction was from the west, therefore trap plants placed at the distances of 50– 75 m east of the infected plot were expected to be uniformly infected). The reasons for this dispersal pattern are not clear, and our analysis is limited by rather limited data. Detailed studies are needed to get a more complete picture of spatial patterns of conidial spread during sprinkler irrigation events.

Above canopy irrigation reduces mean ambient air temperature by 7–9 °C and increases humidity up to 30 % during the irrigation period (Rotem and Palti 1969). This has implications for Ascochyta blight development within an irrigated chickpea paddock, as *A. rabiei* only requires 3–10 h leaf wetness for host penetration and infection at the optimum temperature (Khan 1999; Moore et al. 2016). Therefore, extended periods of sprinkler irrigation events may not be required for disease development, especially in a cool and moist season. In the laboratory; more than 50 % of conidia, incubated on membrane filters, germinated after four days at relative humidity as low as 12.5 % (temperature 20 °C) (Coventry 2012). *Ascochyta rabiei* conidia have been shown to survive and remain infective, when inoculated on chickpea seedlings in a growth chamber, during dry periods lasting 24 h (Armstrong-Cho et al. 2004). However, conidia are unlikely to withstand dry conditions for extended periods of time under field conditions, especially at sub-optimal temperatures. For conidial spread with sprinkler irrigation, this implies that even though conidia may spread to equal distances (75 m in this case) during sprinkler irrigation and rainfall events, leaf wetness provided by dew and/or humidity over 90% to facilitate conidial penetration and infection will be important in establishing disease on chickpea outside the irrigated paddock in the absence of rain (if temperature requirement is met). Conversely; rainfall usually occurs over a large area, and only a few mm of rain later in the day or at night satisfies the leaf wetness requirement (Moore et al. 2016). Therefore, leaf wetness provided by dew and/or high relative humidity is not necessary to facilitate conidial penetration and infection at longer distances for conidia dispersed in rainfall events. This further implies that rainfall might be more important than sprinkler irrigation for conidial spread and subsequent disease development at farther distances. Future work determining the viability of conidia after exposure to different duration of dry periods, under field conditions, is necessary to understand the duration of dry period over which *A. rabiei* conidia can survive when dispersed outside the irrigated paddock. To the best of our knowledge, this is the first systematic study to determine conidia dispersal with sprinkler irrigation.

The finding that conidia travelled 75 m during all rainfall and sprinkler irrigation events suggests that conidia can potentially play a role in the long-distance dispersal of *A. rabiei*. Conidia dispersed 50 m from naturally infested stubble from the previous year (there were no trap plants farther than 50 m to test long-distance dispersal) (Galloway, unpublished). The sexual spores of the fungus (ascospores) have been reported to travel up to hundreds of metres to several kilometres from the source of infection (Trapero-Casas et al. 1996; W. Kaiser and Küsmenoglu 1997; Trapero-Casas and Kaiser 1992). The present study did not have trap plants farther than 75 m from the infected plot and wind speed during spread events was rather low with a maximum mean wind speed of 6.5 ms^-1^ and maximum gust of 11.2 ms^-1^. It is likely that conidia could dispersed farther than 75 m, especially when wind speed is high and there are strong wind gusts. It has been hypothesized that if conidial dispersal is aided by turbulent winds and conidia enter the turbulent boundary layer, they could be dispersed to distances that are associated with ascospores only (McCartney and West 2007; Coventry 2012). While the role of conidia in long-distance dispersal has not been recognized, it has been hypothesized that liquid suspensions in air (aerosols) can incorporate a large number of conidia in heavy rain that can be transported over longer distances by wind (Coventry 2012). A severe Ascochyta blight epidemic developed across the southern region of Australia in 1998. It is speculated that aerosols developed during heavy rains, which travelled over hundreds of meters to kilometers and contributed to the epidemic development (Coventry 2012). This suggests that role of conidia in the long-distance dispersal has been underestimated. Further work distributing trap plants at distances farther than 75 m is necessary to determine the potential distance conidia can travel.

When fitting observed data to logistic models, the known biology of pathogens needs to be considered rather than simply considering the shape of the curve (Sparks et al. 2008). Generalised additive models are data-driven not model-driven, as they allow the data to determine the shape of the response curves rather than being limited by the shapes attainable through some parametric specification. The inability to control environmental factors, such as wind speed, amount of rainfall, temperature and humidity makes it difficult to use parametric statistical models e.g., generalised linear models (GLMs) to make inferential comparisons across sites without a high risk of the presence of confounding variables.

Generalised additive models offer a flexible modelling approach that provides a good balance between highly interpretable linear models that have undesirable predictive capabilities and black-box machine learning approaches, such as neural networks, which have excellent predictive performance but low interpretability (Hastie and Tibsbirani 1990). This capacity to achieve interpretable separation of covariate contributions in a strong descriptive model is the main reason that GAMs were chosen to model spatial patterns of conidial spread. A model was developed that efficiently described spatial patterns of conidial spread under field conditions. The model will be useful for chickpea growers in identifying the scale of potential damage and could be used to predict the spread of *A. rabiei* with further development. When further developed, this model can also be used to inform chickpea growers how far to grow chickpea from the neighbouring paddocks.

A comparison of conidial spatial dispersal patterns in identical amount of irrigation and rainfall events would have been beneficial, but constantly changing wind speed and direction makes a direct comparison difficult under field conditions. Conidial spread was investigated in the late stage of plant growth with 90–100 % plants in plots being infected. Detailed studies at different plant growth stages and disease incidence levels are now needed to provide a more complete picture of distance conidia disperse and their spatial dispersal patterns. It should be noted that counting lesions on trap plants gave an estimate of only viable conidia dispersed (not the total number of conidia dispersed), which could cause disease under field conditions if conditions are favourable for disease development. This study was primarily conducted to get an estimate of the distance conidia travel and remain viable. Future studies should consider using trap plants, spore traps and molecular approaches (e.g. trapping spores sticky tapes and subsequent spore count using quantitative PCR) to get an accurate estimate on the number of conidia dispersed. Spore trapping coupled with quantitative PCR has been demonstrated to be a useful tool for epidemiological studies and biosecurity surveillance (Vogelzang 2012). More work is required to establish the equivalency of number of lesions to the number of conidia. Many other factors are involved in the spore dispersal, including changing wind directions, turbulence and topographic factors (Coventry 2012), which add complexity to the model and were not considered in the present model. These factors need to be addressed in future models.

## Acknowledgements

Grains Research and Development Corporation (GRDC), Australia, provided financial assistance in this work through the project USQ-1903-003RTX. Agriculture Victoria and GRDC provided financial assistance in this work through their co-investment project DAV00150, and DJP1097-001RTX.

We thankfully acknowledge Jason Brand and his team for trial sites management. We thank Andrew Hallet for technical support and assistance with field work.

## Ethical approval

This study did not involve working with animals or humans

## Consent to participate

Not applicable

## Consent for publication

Not applicable

## Availability of data and material

The raw data are documented and available from https://doi.org/10.5281/zenodo.3842293.

All raw and generated data and further associated materials have been made further available as a part of a research compendium available from https://doi.org/10.5281/zenodo.3810826.

## Code availability

All code used in the analyses and data visualisation have been made available as a research compendium, available from https://doi.org/10.5281/zenodo.3810826.

---

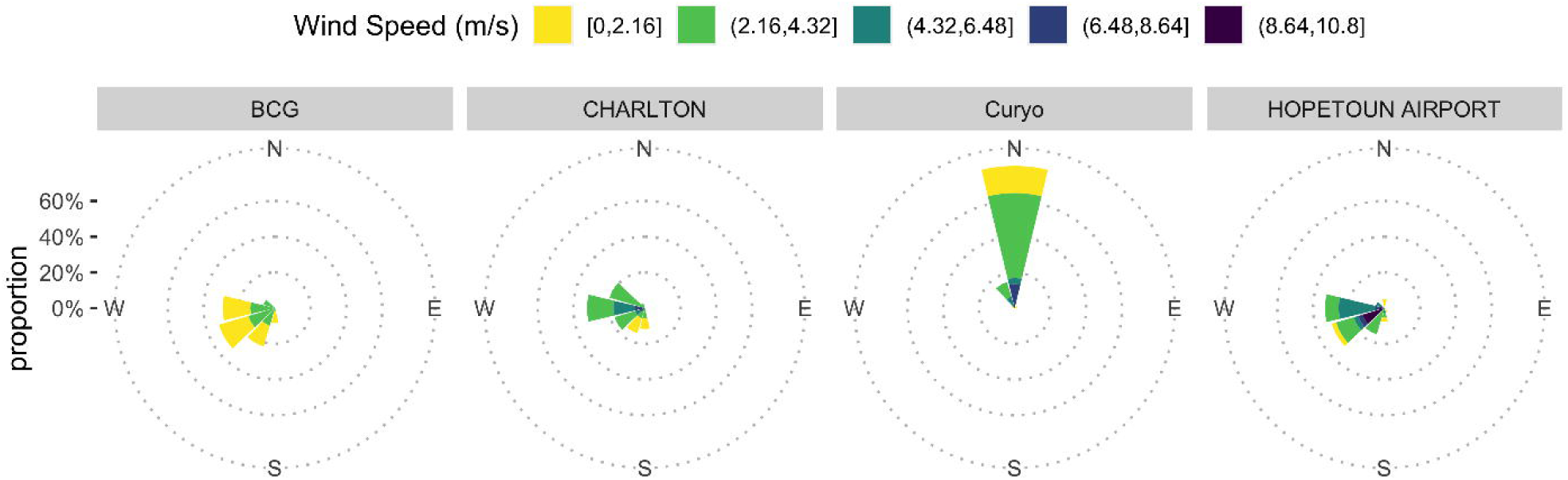

